# Quantile normalization of single-cell RNA-seq read counts without unique molecular identifiers

**DOI:** 10.1101/817031

**Authors:** F. William Townes, Rafael A. Irizarry

**Affiliations:** Department of Computer Science, Princeton University, Princeton, NJ; Department of Data Sciences, Dana-Farber Cancer Institute, Boston, MA; Department of Biostatistics, Harvard University, Cambridge, MA

**Keywords:** gene expression, single cell, RNA-seq, normalization, quasi-UMI

## Abstract

Single-cell RNA-seq (scRNA-seq) profiles gene expression of individual cells. Unique molecular identifiers (UMIs) remove duplicates in read counts resulting from polymerase chain reaction, a major source of noise. For scRNA-seq data lacking UMIs, we propose quasi-UMIs: quantile normalization of read counts to a compound Poisson distribution empirically derived from UMI datasets. When applied to ground-truth datasets having both reads and UMIs, quasi-UMI normalization has higher accuracy than alternatives such as census counts. Using quasi-UMIs enables methods designed specifically for UMI data to be applied to non-UMI scRNA-seq datasets.

## 1 Background

Single cell RNA-seq (scRNA-seq) has become a standard tool for measuring gene expression patterns from individual cells. The initial molecule capture and reverse transcription (RT) steps in scRNA-seq protocols result in low quantities of cDNA, so a large number of PCR cycles are needed to produce enough material for measurement. The resulting libraries, that are then sequenced, contain many duplicates of each of the single mRNA molecules extracted from the original cell [1]. To account for this distortion, current protocols include unique molecular identifiers (UMIs), which enable computational removal of PCR duplicates [2]. However, read count datasets generated without UMIs are still in use for at least two reasons. The first is that many public datasets have been produced with non-UMI protocols. The second is that current UMI protocols sequence only the 5-prime or 3-prime end of the mRNA molecule, and therefore prevent quantification of transcript isoform levels within the same gene [3] or allele specific expression [4].

In both UMI and read count data, the fraction of zeros per cell is often a dominant source of variation. Not only does the zero fraction strongly correlate with the first principal component, but it also affects the entire gene expression distribution [5]. This is a result of cell-to-cell differences in capture and RT efficiency, which has nothing to do with underlying biology. For UMI counts, systematic variation introduced by these technical components can be addressed by using multinomial models [6]. However, for read counts, such models are precluded by the additional multiplicative distortions of PCR. Here, we focus on the analysis of read counts from non-UMI protocols such as Smart-seq2 [7]. Note however that read counts (with PCR bias) may also be obtained from UMI protocols if the UMIs are simply ignored when constructing the expression measurements.

The substantial distortions in read counts have motivated the development of sophisticated normalization procedures. One approach is to attempt to transform the data to more closely follow a normal (Gaussian) noise model. For example, log-normalized counts from transcripts per million (TPM), SCRAN [8], or SCnorm [9] may be used as input to principal component analysis (PCA) which implicitly assumes Gaussian noise. However, due to the large number of zeros in scRNA-seq, log transformation of normalized counts requires a pseudo-count, which introduces substantial bias [10]. The resulting distributions can be far from Gaussian, even for UMI count data [6]. In contrast, the *census counts* method transforms read counts and attempts to match the underlying UMI distribution based on the key observation that the mode of the nonzero UMI count distribution is typically one. Rather than matching a normal distribution, this approach needs only to remove PCR bias to be effective. The resulting census counts can be analyzed as if they were UMI counts by methods specifically developed for UMI data [11, 6]. Census count normalization relies on a complex mechanistic model of scRNA-seq biochemistry and applies a linear transformation [1]. However, due to the nonlinearity of PCR, this approach is inadequate for removing bias.

Here we present quasi-UMIs (QUMIs), a normalization technique for scRNA-seq read counts that, like census counts, attempts to match the UMI count distribution. Our approach differs from census counts in that we apply quantile normalization rather than a linear transformation, producing a discrete distribution. In general, quantile normalization forces all cells to follow a specific *target distribution*. The most widely implemented version of quantile normalization generates the target distribution by averaging over empirical distributions from the data [12]. In the case of scRNA-seq, however, we know that if we could remove PCR duplicates from read counts we would obtain UMI counts, which have a markedly different distribution from any of the empirical read count distributions [6]. We therefore use the characteristics of UMI counts as a guide to construct the QUMI target distribution such that it will approximate a true UMI count distribution. Specifically, we fit Poisson-lognormal models to three public datasets from different species and UMI protocols. The Poisson-lognormal distribution has a heavy tail that approximates a power law, and power laws have been observed previously in gene expression [13, 14]. This target QUMI distribution depends on a single shape parameter and makes no assumptions about biochemical mechanisms. On three independent benchmark datasets where both read and UMI counts were available, we transformed read counts to QUMIs and census counts. We assessed accuracy by computing distances between normalized read counts and the true UMI counts. QUMIs had higher accuracy than census counts and read counts. Finally, on a dataset without UMIs, using QUMIs combined with a UMI count based dimension reduction reduced batch effects and increased detection of biological groups.

## 2 Results and discussion

### 2.1 Datasets

We used seven public scRNA-seq datasets (Table 1). For the three training datasets, we only obtained UMI counts. For the three test datasets, we obtained both UMI counts and read counts. Finally, for the prediction dataset, we obtained only read counts. We refer to each dataset using the first author’s last name.

**Table 1:**
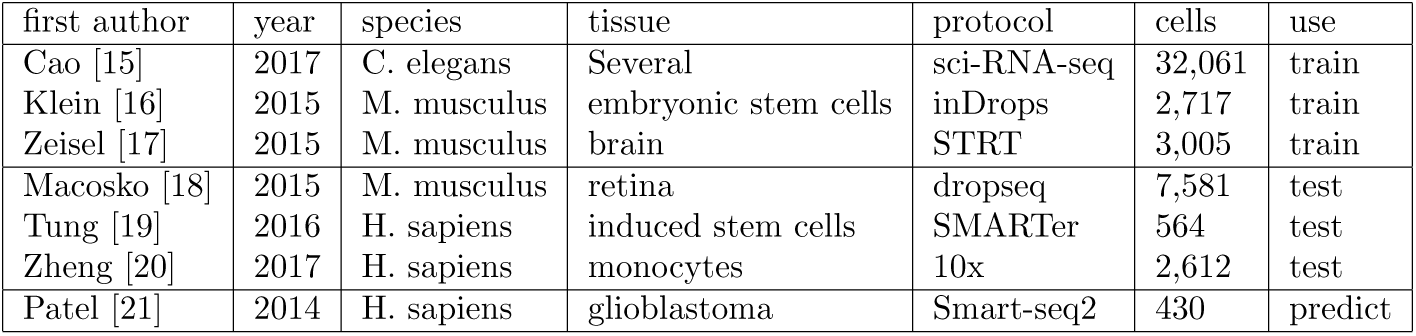
Single cell RNA-seq datasets used. Training data contained only UMI counts. Test data contained UMI counts and read counts. Prediction data contained only read counts.

| first author | year | species | tissue | protocol | cells | use |
| --- | --- | --- | --- | --- | --- | --- |
| Cao [15] | 2017 | C. elegans | Several | sci-RNA-seq | 32,061 | train |
| Klein [16] | 2015 | M. musculus | embryonic stem cells | inDrops | 2,717 | train |
| Zeisel [17] | 2015 | M. musculus | brain | STRT | 3,005 | train |
| Macosko [18] | 2015 | M. musculus | retina | dropseq | 7,581 | test |
| Tung [19] | 2016 | H. sapiens | induced stem cells | SMARTer | 564 | test |
| Zheng [20] | 2017 | H. sapiens | monocytes | 10x | 2,612 | test |
| Patel [21] | 2014 | H. sapiens | glioblastoma | Smart-seq2 | 430 | predict |

### 2.2 Current normalization methods inadequate for scRNA-seq read counts

We obtained read counts from UMI protocols and produced both UMI counts and, by ignoring the UMIs, read counts. For each gene we then had both UMI and read count measurements. The variability introduced by PCR resulted in hundreds of genes with read counts above 100 that mapped back to less than five UMI counts, in some cases just one (Figure 1). Current normalization methods such as transcripts or counts per million (TPM, CPM) and census counts apply linear transformations to read counts from non-UMI protocols, which preserve the PCR distortions and result in variable distributions even when the data are generated with the same cell type [19] (Figure 2a-c). Different distributions can be observed when data are processed in different batches (Figure 2d-f) which can then lead to apparent differences in the low-dimensional representations used, for example, to discover new cell types [5]. For example, the Patel dataset [21] consists of five glioblastoma tumors, with one of these processed in two batches. Current normalizations do not remove the substantial variation in distributions between the batches (Figure 2e-f). Since not only the scale but also the shape of the distribution of expression values is highly variable between cells of the same biological condition, normalization based on linear transformation is insufficient.

**Figure 1:**
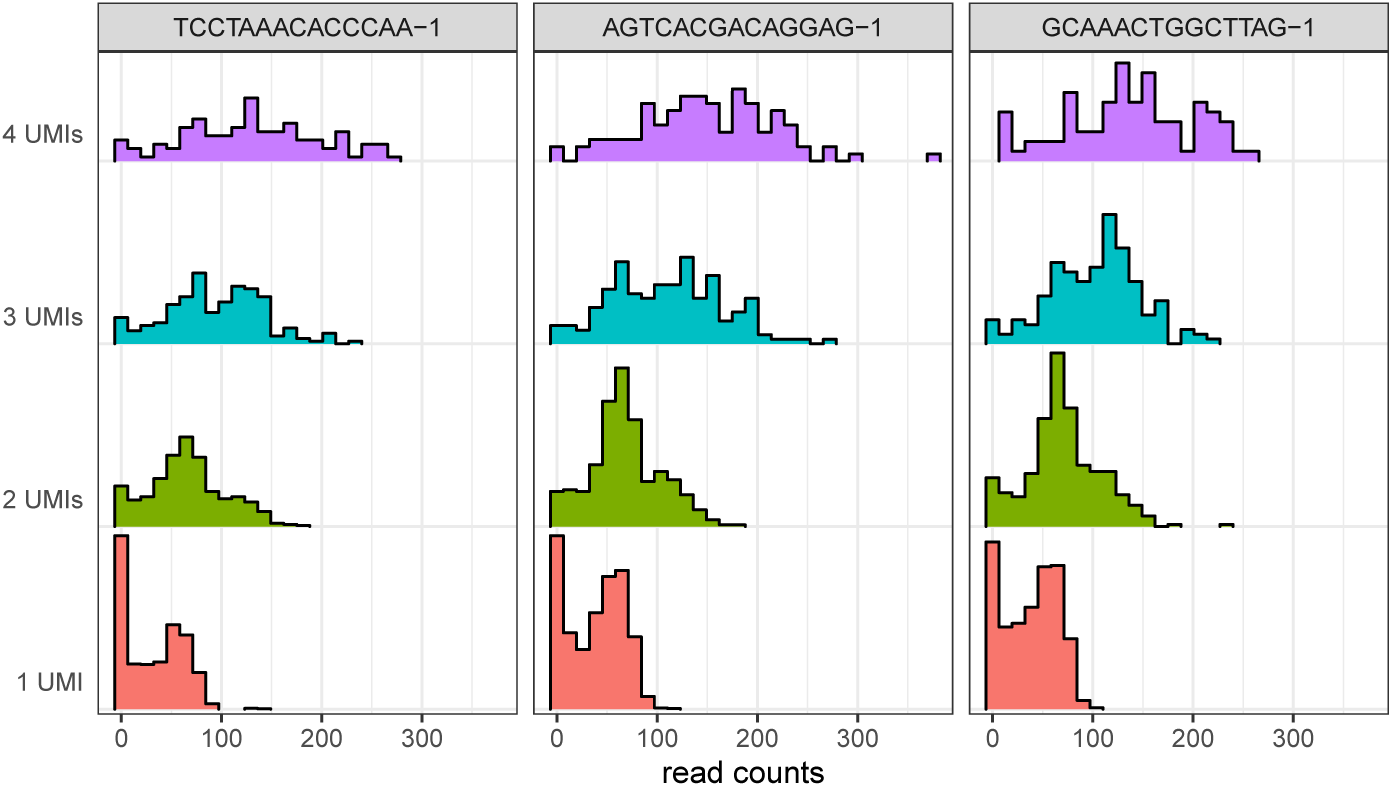
PCR amplification produces a wide range of read count values originating from genes represented by small UMI count values in the Zheng dataset. Each pane is a cell. Color indicates UMI count value.

**Figure 2:**
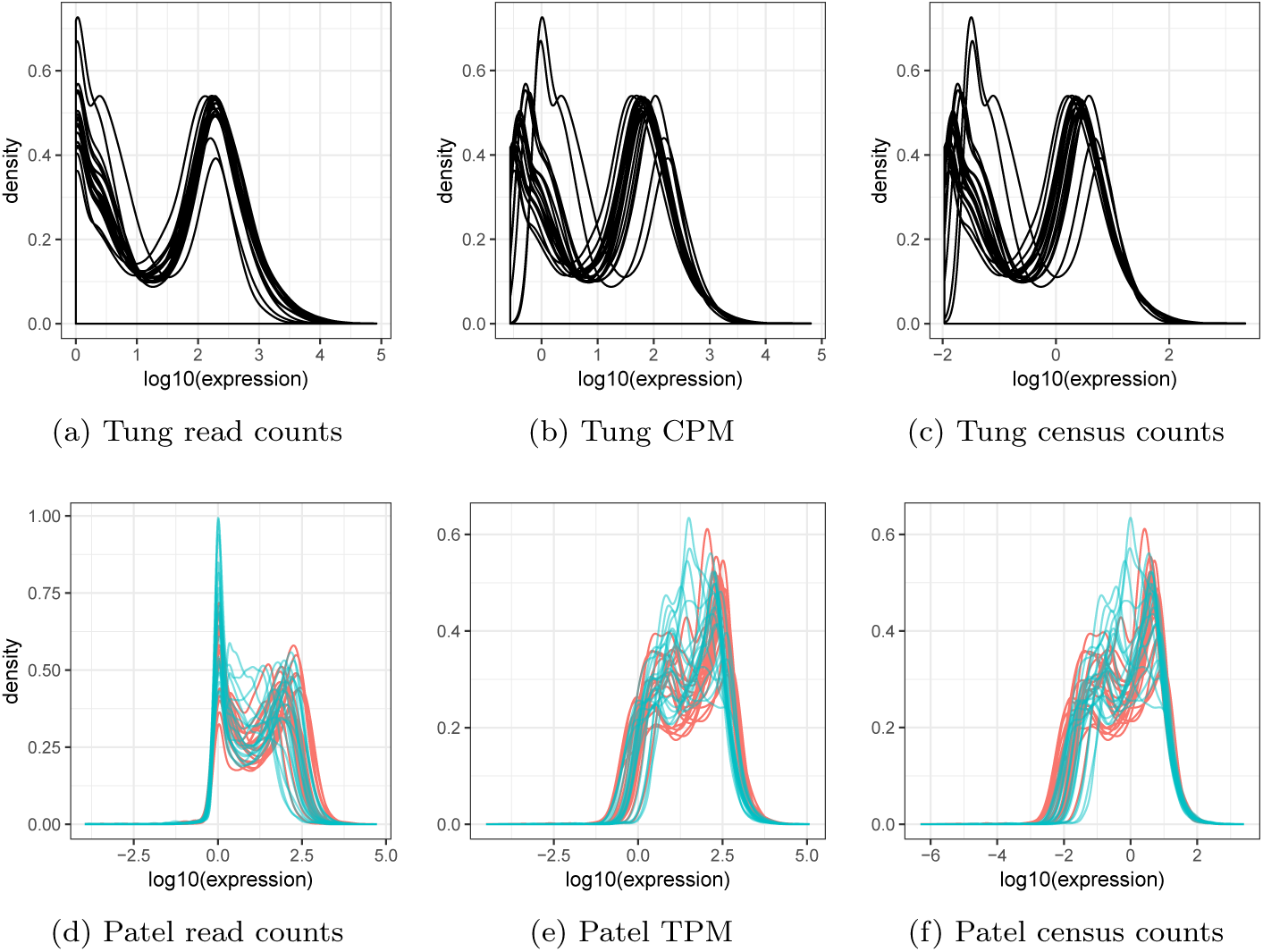
Current normalizations do not remove distributional variation across biological replicates and batches. a) Kernel densities of nonzero read count values for 20 random cells from individual NA19098, replicate r3 of the Tung dataset. b) as a) but normalized to counts per million. c) as a) but normalized to census counts. d) Kernel densities of nonzero read count values for 20 random cells from each batch of tumor MGH26 in the Patel dataset. Color indicates batch. e) as d) but normalized to transcripts per million. f) as d) but normalized to census counts.

### 2.3 UMI counts fit by Poisson-lognormal distribution

To quantile normalize read counts to match UMI counts, we identified the qualitative characteristics of the target UMI count distribution. Due to the heavy tail of UMI counts, log-log plots are an effective way to visualize their distribution (Figure 3a). Log-log plots are essentially histograms with both axes log transformed and if the right tail of the distribution appears linear it is suggestive of a power law distribution [22]. Stacking log-log plots for 500 randomly chosen cells, we observed a monotonic decreasing trend for all cells but with substantial variability in the observed proportions for each UMI count value (Figure 3b). Consistent with [1], the most prevalent nonzero value was one.

**Figure 3:**
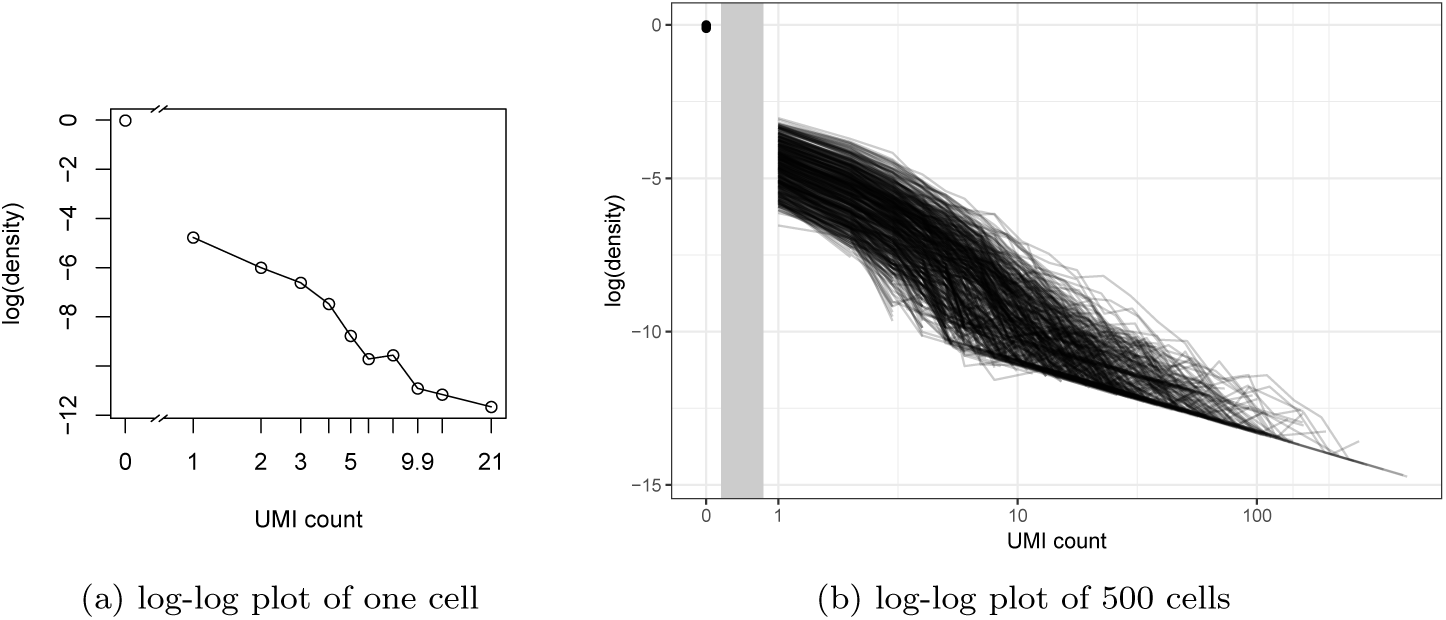
Log-log plots reveal monotonically decreasing, heavy tailed distributions in UMI counts. a) Log-log plot of UMI counts from cell cele-002-090.AATCATACGG in the Cao dataset. b) as a) but with 500 random cells. Vertical gray bar indicates discontinuity due to horizontal axis log scaling.

A recent survey of a wide variety of datasets found that ostensible power law relationships are better described by lognormal distributions [23]. We therefore considered both the Poisson-Lomax, which has a true power law tail, and the Poisson-lognormal families as candidates for the quantile normalization target distribution. Probability mass functions (PMFs) are listed in Methods. While both distributions fit the training data well overall, the Poisson-Lomax tended to overestimate the probability of high magnitude outliers (Figure 4). Also, maximum likelihood estimation of the parameters of the Poisson-Lomax model was numerically less stable. Hence, we focused on the Poisson-lognormal model in our subsequent assessments.

**Figure 4:**
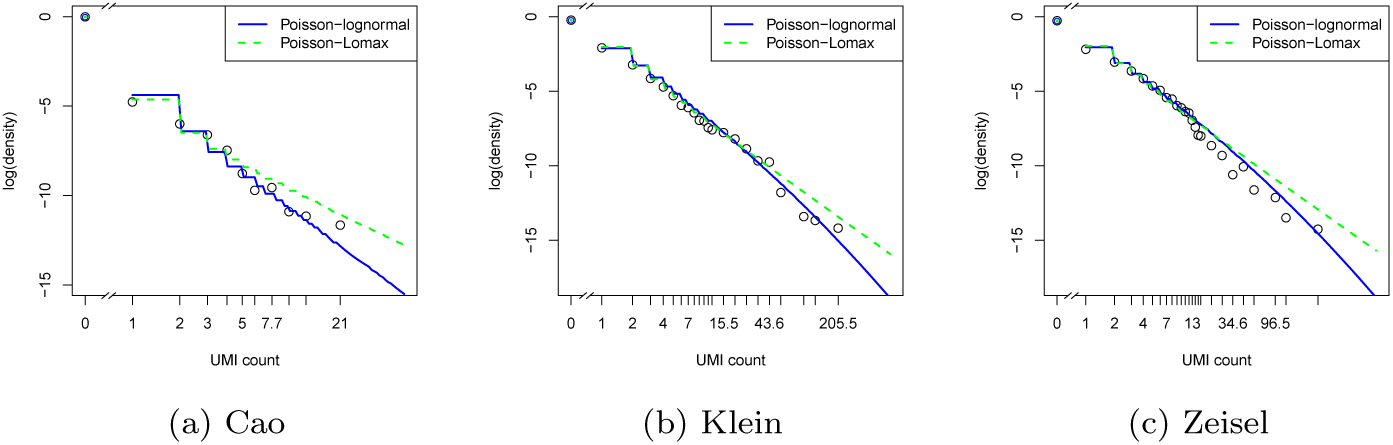
Log-log plots of UMI counts (points) with maximum likelihood fits (curves). a) Cell cele-002-090.AATCATACGG in the Cao dataset. b) Cell 1146 from the d2 group in the Klein dataset. c) Cell 1772067059 H11 from the Zeisel dataset.

### 2.4 Quantile normalization of read counts to quasi-UMIs

Assuming the underlying UMI count distribution is Poisson-lognormal, only two parameters are needed to describe each cell: scale and shape. If UMI data is available these are easily estimated using maximum likelihood (MLEs). However, in read count data without UMIs this is not possible due to PCR distortion. Conveniently, if the shape parameter is known a priori, the scale parameter can be estimated from the fraction of zeros. This is useful because the zero fraction derived from read counts equals the zero fraction in the UMI counts for the same cell; zero is the only expression value in read count data that is not altered by PCR bias. Therefore, if the shape parameter is assumed known, the target distribution for a given cell can be determined from the read count data. Our method requires the shape parameter to be fixed and to determine reasonable default shape parameter values, we computed MLEs for all cells in the training data (Figure 5).

**Figure 5:**
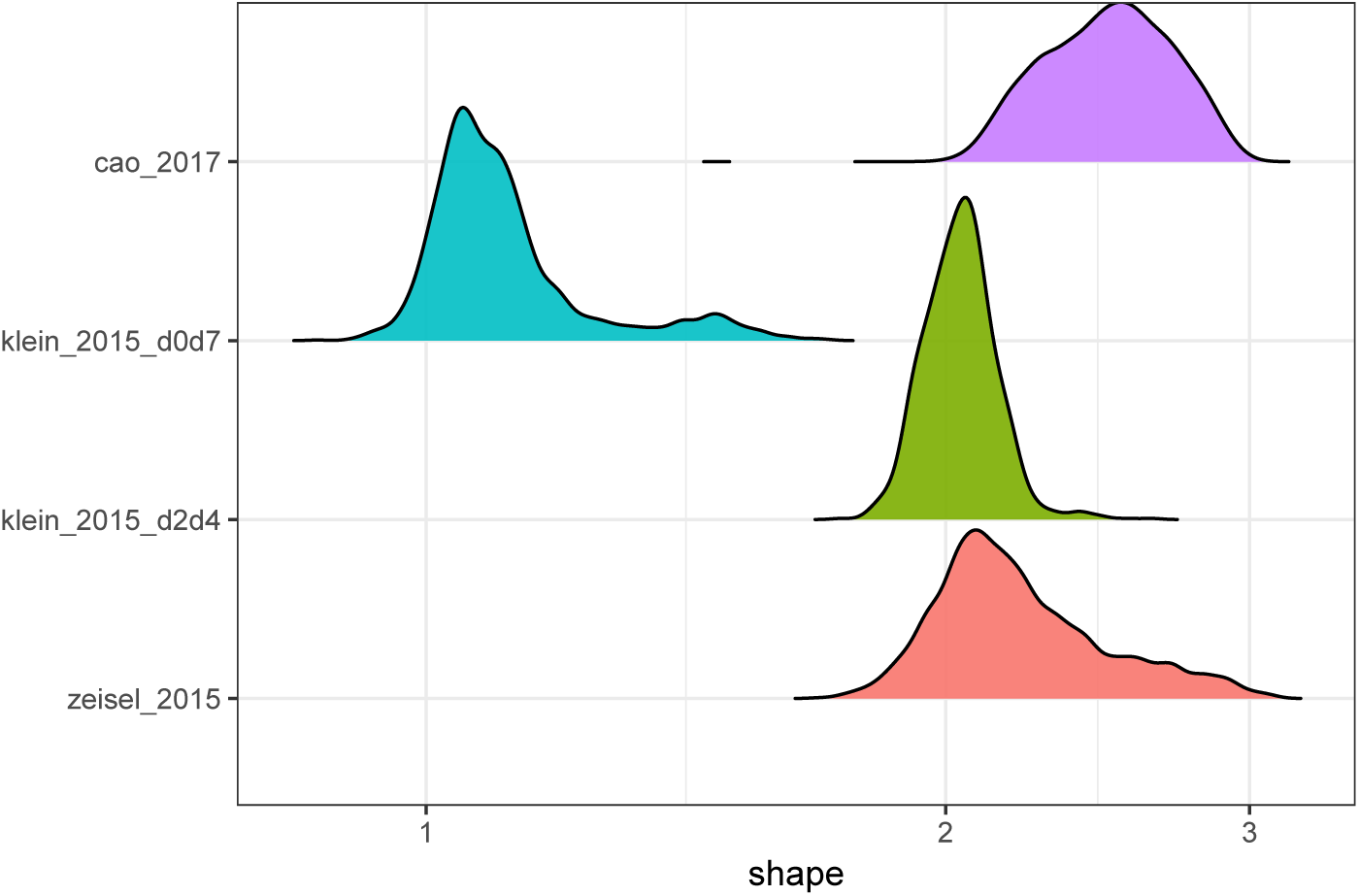
Poisson-lognormal shape parameter maximum likelihood estimates (MLEs) for the three training datasets. klein 2015 d0d7 and klein 2015 d2d4 indicate different experimental conditions within the same dataset.

While it was necessary to fix the shape parameter before applying quantile normalization to the test data, each cell was allowed to have its own scale parameter. If the scale parameter were also held fixed, the QUMI target distribution would be identical for every cell and would predict a constant zero fraction across cells. But this is discordant with the fact that UMI count data exhibit variation in the zero fraction across cells [6]. Since the varying zero fractions in read counts exactly match the zero fractions in underlying UMI counts, it would be inappropriate to alter these correct expression values by normalizing to a global target distribution. Instead, we estimated each cell’s scale parameter directly from the zero fraction in read counts using the method of moments (MOM). A detailed explanation of the estimation procedure is provided in the Methods. Because this approach matched each cell’s zero pattern, only the nonzero read counts needed to be adjusted by the normalization, which improved computational efficiency.

After estimating the scale parameter for each cell, we obtained empirical quantiles (ranks) from read counts, and transformed the ranks to QUMI counts by matching to the target distribution’s theoretical quantiles (see Methods for detailed algorithm).

### 2.5 Quasi-UMIs approximate UMIs more closely than census counts

Using the three test dataset UMI counts as ground truth, we compared the accuracy of Poisson-lognormal quasi-UMI (QUMI) counts with census counts and unnormalized read counts. We quantified the accuracy of a normalization method for a given cell by computing the Euclidean distance between the log of the normalized count vector and the log of the UMI count vector. Zero values were omitted from the computation because all of the normalization methods preserved the sparsity structure of the read counts.

Across datasets, QUMI counts had the highest accuracy (smallest median distance from UMIs counts), while census counts were more accurate than read counts (Figure 6). The improvement from using QUMI normalization was most dramatic on the deeply sequenced Tung dataset. The Macosko and Zheng datasets came from droplet protocols with shallow sequencing, while the Tung data came from a plate protocol. The latter is more similar to non-UMI protocols such as Smart-seq2, suggesting QUMI normalization is likely to be effective in those settings.

**Figure 6:**
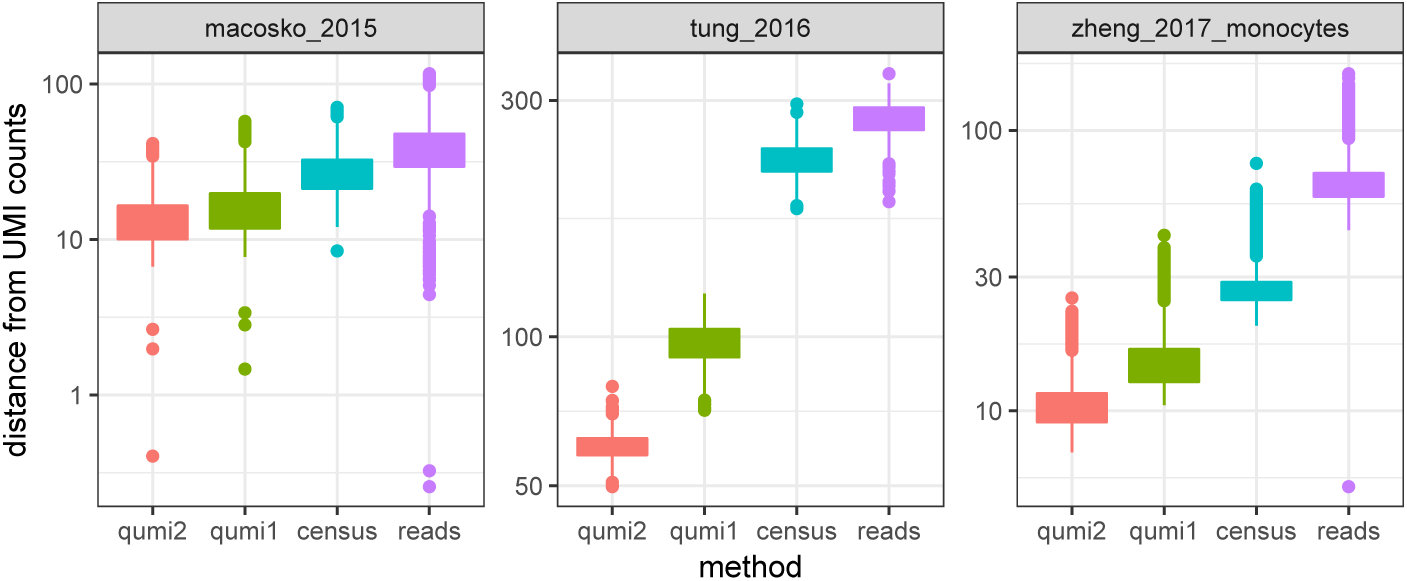
Quasi-UMI counts approximate UMI counts more closely than census counts. qumi1,qumi2: Poisson-lognormal QUMI counts with shape parameters of 1.0 and 2.0 respectively, census: census counts, reads: read counts.

As a sensitivity analysis, we repeated the QUMI normalization for all datasets with fixed shape parameter values of 1.0 and 2.0. While altering the shape parameter affected the accuracy of QUMI counts, with shape of 2.0 leading to better accuracy, the difference was small compared to the difference between QUMIs and census counts (Figure 6).

### 2.6 Quasi-UMIs enable dimension reduction of read counts

Quasi-UMI counts may be analyzed as if they were UMI counts. To illustrate this, we applied the Poisson-lognormal QUMI normalization (with shape 2.0) and census to TPM values from the Patel dataset [21]. We used TPM values instead of raw read counts as input because full-length scRNA-seq protocols exhibit gene length bias [24]. This dataset lacked UMIs and profiled 430 cells from five glioblastoma tumors. One tumor (MGH26) was processed in two batches on two different sequencing machines. These two batches differed in the fraction of zeros [5]. We then reduced the dimensionality with GLM-PCA [6] and compared this to PCA dimension reduction on *log*_2_(1 + *T P M*) from the original read count data. GLM-PCA was specifically designed to handle UMI counts. Only the 5, 685 genes used by the original authors were included as input to dimension reductions.

PCA applied to log-transformed TPM from read counts failed to merge the two batches of MGH26 and poorly separated tumors MGH28, MGH29, and MGH30. In contrast, GLM-PCA applied to either QUMI or census counts merged MGH26 batches and separated all tumors. GLM-PCA applied to QUMI counts better separated MGH31 tumor compared to census counts (Figure 7). Consistent with [5], the dominant source of variation in log-transformed TPM was the fraction of zeros; the correlation between the first PC and zero fraction was 0.96. GLM-PCA reduced correlations between the first latent dimension and zero fraction to 0.58 and 0.26 for census and QUMI counts respectively (Figure 8). This showed that QUMI normalization, like UMIs, does not remove all technical variation. Rather, it enables application of methods specifically designed for analysis of UMI counts to datasets that lack UMIs.

**Figure 7:**
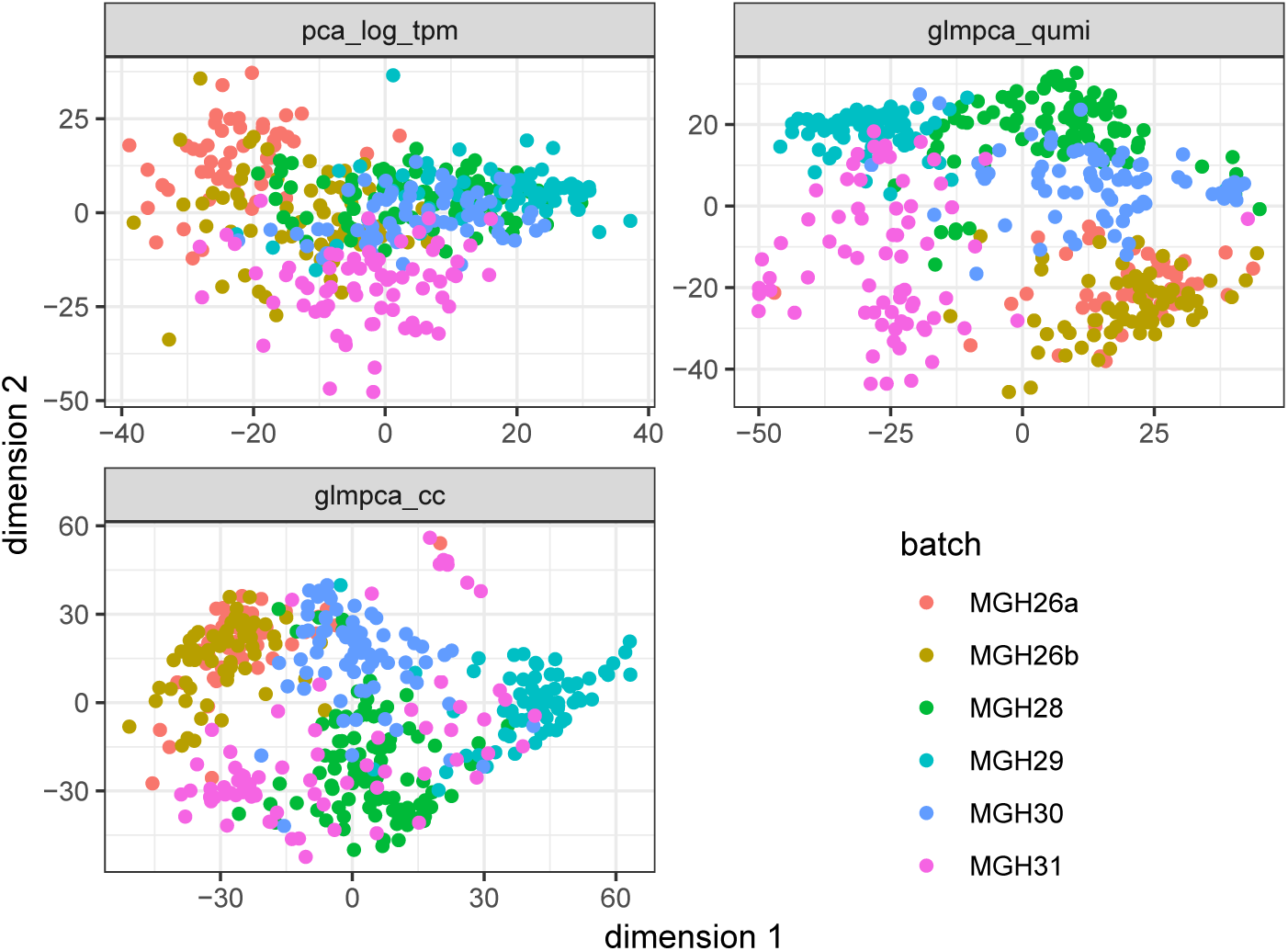
Dimension reduction of non-UMI glioblastoma dataset before and after QUMI and Census normalizations. pca_log_tpm: PCA on transcripts per million from original data, glmpca_cc: GLM-PCA on census counts, glmpca_qumi: GLM-PCA on QUMI counts.

**Figure 8:**
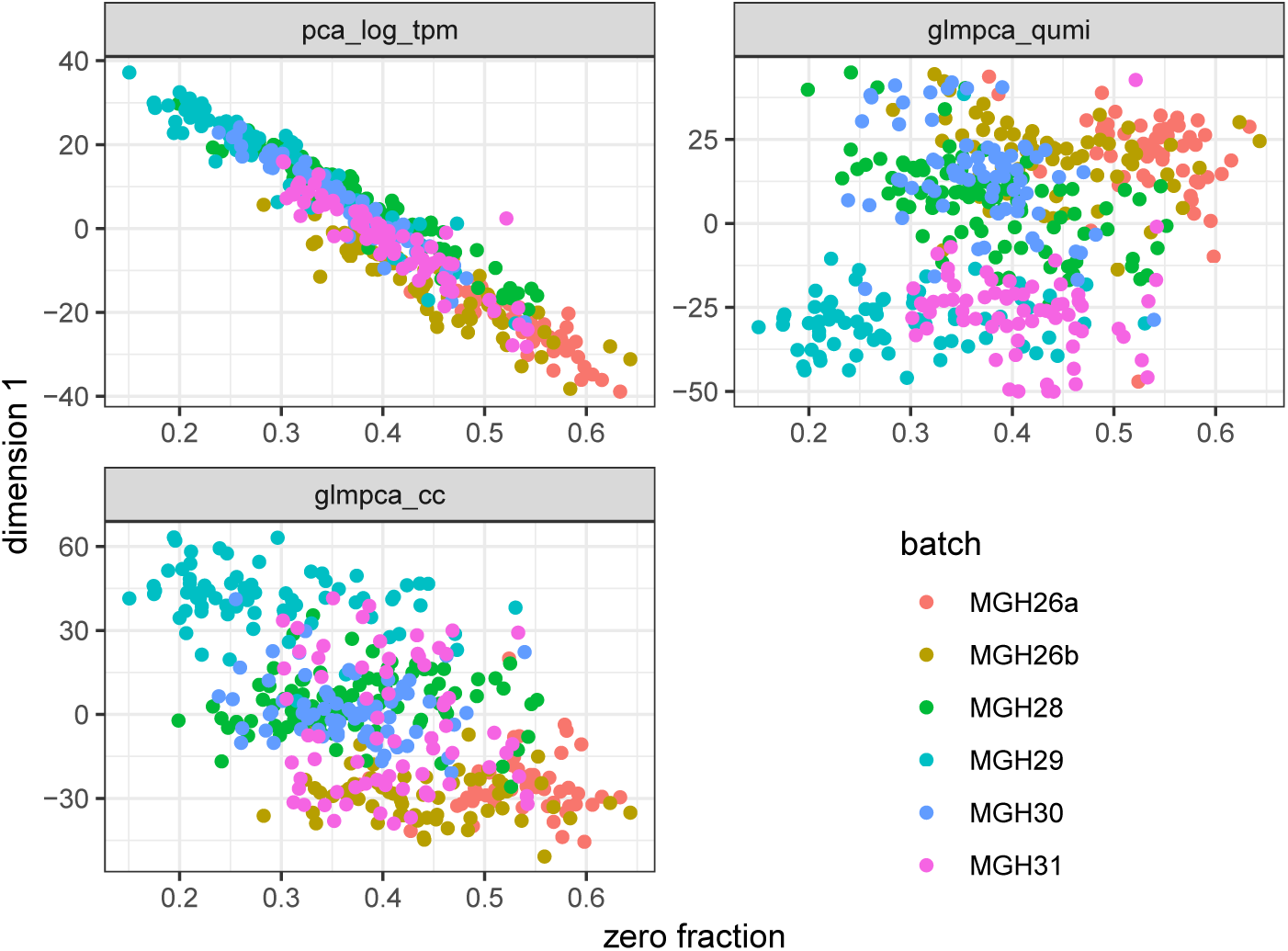
GLM-PCA reduces correlation between zero fraction and first latent dimension in a non-UMI glioblastoma dataset. pca log tpm: PCA on transcripts per million from original data, glmpca cc: GLM-PCA on census counts, glmpca qumi: GLM-PCA on QUMI counts.

## 3 Conclusion

We have shown that UMI counts can be approximated by quantile normalization of read counts to quasi-UMIs (QUMIs) in scRNA-seq. The Poisson-lognormal model fits UMI count data well and can be used as a target distribution for QUMI normalization. However, the conceptual framework is generalizable to any discrete distribution that can be calibrated against UMI data, such as the Poisson-Lomax or two-component mixture models of active and inactive genes. Using test datasets with read counts and UMI counts from the same cells, we confirmed QUMI counts approximate UMI counts more closely than census counts and unnormalized read counts.

QUMI normalization mitigates the distortion of PCR amplification in scRNA-seq protocols that lack UMIs while preserving sparsity. However, just like the use of proper UMIs, it does not normalize differences between cells resulting from variation in efficiency of capture or reverse transcription. This contributes to differences in the zero fraction across cells, which are intentionally preserved in QUMI normalization. These sources of technical variation should be addressed through UMI-specific count models such as GLM-PCA or its approximations using residuals [6, 11].

QUMI counts do not directly account for PCR bias arising from differences in gene length or GC content. These biases are not specific to single cell protocols and have been addressed in the bulk RNA-seq literature [25]. Since QUMI normalization only requires the rank ordering of genes in each cell along with the fraction of zeros, bias-adjusted TPM values from pseudoaligners [26, 27] can be used as input instead of raw read counts. We followed this approach in analyzing the Patel data.

A major advantage of QUMI counts is that they can be analyzed as if they were UMI counts. This avoids the need to develop customized methods of dimension reduction and feature selection for the read count distribution. Here, we have focused specifically on scRNA-seq read counts, but traditional bulk RNA-seq read count data is also affected by distortion from PCR amplification. While it may be possible to extend the QUMI framework to bulk RNA-seq data, an appropriate target distribution would need to be identified. This is challenging because a bulk RNA-seq sample, unlike scRNA-seq, is typically a mixture of cell types with unknown proportions. Such a mixture is unlikely to be easily characterized by a simple two parameter distribution. PCR distortion is also present in read counts from metagenomics experiments [28].

Finally, we caution that QUMIs are not substitutes for proper UMIs. If the latter can be used in an experiment, they will certainly be more effective than QUMIs in removing PCR distortions. QUMI normalization relies on assumptions, such as a global, fixed shape parameter, which may not be met in certain datasets. Indeed, we observed in the Klein dataset that the shape parameter was not constant across experimental conditions, although our method was robust to this choice.

## 4 Methods

### 4.1 Data acquisition and preprocessing

The Cao dataset UMI counts [15] were obtained by following instructions on the authors’ website http://atlas.gs.washington.edu/worm-rna/docs/. A preprocessed version of the Klein dataset UMI counts [16] was downloaded from https://hemberg-lab.github.io/scRNA.seq.datasets/mouse/esc/. The Zeisel dataset UMI counts [17] were downloaded from the authors’ website http://linnarssonlab.org/cortex/ and low quality cells were removed according to the same criteria used in the original publication. Read and UMI counts for the Zheng dataset [20] were obtained by processing the permolecule information file from https://support.10xgenomics.com/single-cell-gene-expression/datasets/1.1.0/cd14_monocytes. Read and UMI counts for the Macosko dataset [18] were obtained by pseudoaligning raw FASTQ files from Sequence Read Archive using Kallisto version 0.45.1 [26] to produce BUS files [29]. We only included sample r6 from this dataset. Read and UMI counts for the Tung dataset [19] were obtained by following instructions on the authors’ website https://jdblischak.github.io/singleCellSeq/analysis/compare-reads-v-molecules.html. Finally, the glioblastoma dataset read counts and TPM values [21] were downloaded from https://github.com/willtownes/patel2014gliohuman.

### 4.2 Compound Poisson distributions

The probability mass function (PMF) of a compound Poisson distribution is obtained by placing a prior on the rate parameter of an ordinary Poisson distribution.

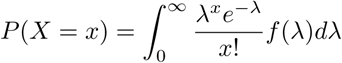

For the Poisson-lognormal distribution with shape (logarithmic standard deviation) *σ* and scale (logarithmic mean) *µ*, the prior is a lognormal distribution with the same parameters:

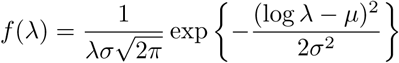

For the Poisson-Lomax distribution with shape (power law tail index) *α* and scale *θ*, the prior is a Lomax (shifted Pareto) distribution with the same parameters:

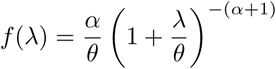

Let *m* and *v* represent the mean and variance of a given prior distribution in a compound Poisson model. The marginal mean of the compound Poisson is also *m* and the marginal variance is *m* + *v*. For example, for the Poisson-lognormal, the mean is exp(*µ* + *σ*^2^*/*2) and the variance is *m* + exp(*σ*^2^) *−*1 *m*^2^. The Poisson-Lomax distribution has such a heavy tail that its moments are only finite in certain parameter regions. If *α >* 1 then the mean is *θ/*(*α −* 1). If *α >* 2 then the variance is 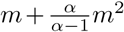. The quadratic variance function in both families is shared with the negative binomial distribution, so none of them can be distinguished based on coefficient of variation. The Poisson-lognormal has a strictly heavier tail than negative binomial, and Poisson-Lomax has a strictly heavier tail than Poisson-lognormal.

For Poisson-lognormal we evaluated the PMF using the R package **sads**. For Poisson-Lomax we evaluated the PMF by using 1, 000 numerical quadrature points. For each cell in the training datasets, we obtained maximum likelihood estimates (MLEs) of compound Poisson model parameters (shape, scale) by numerical optimization using the R function **optim**. The mode of the shape parameter distribution across cells was then used to calibrate the quasi-UMI target distribution in the test and prediction datasets.

### 4.3 Computing quasi-UMIs from read counts

#### 4.3.1 Method of moments estimates from zero fractions

For each cell in the test data, we obtained a target quasi-UMI distribution by estimating the cell-specific scale parameter from the empirical zero fraction in read counts using the method of moments (MOM). Specifically, let *f* (*x*; *µ_i_*) be the Poisson-lognormal probability mass function (PMF) with fixed shape parameter *σ* and unknown cell-specific scale parameter *µ_i_*. For a given cell *i*, the theoretical probability of a zero is *f* (0; *µ_i_*) (a function of *µ_i_* only). The empirical probability of zero is simply the fraction of genes with zero read counts in that cell, which we denote with 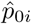. A MOM estimate of *µ_i_* is obtained by finding a root of the function 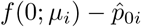 with respect to *µ_i_*.

#### 4.3.2 Quantile normalization

Once a target distribution for an individual cell was determined, we computed the log of the theoretical CDF by cumulatively applying the log-sum-exp transformation to the log-PMF function, which provided numerical stability. We then renormalized the probability distribution to exclude the zero value since zero values in read counts result from UMI counts of zero and do not need to be adjusted. This resulted in a table with positive integer indices providing the quasi-UMI count value and corresponding zero-truncated CDF values indicating the probability of a random variable with the target distribution falling below that value, conditional on it being nonzero. We then converted the vector of read counts from all genes in the cell to empirical quantiles (ranks). Each gene was then aligned to a CDF bin based on its rank. For example, if the zero-truncated CDF had values of 0.8 at 1 and 0.9 at 2, the first 80% of genes with lowest nonzero read count values would be assigned QUMI value of 1 and the next lowest 10% of genes would be assigned QUMI value of 2. Typically a single gene was placed into the highest QUMI bin due to the heavy tail of the target distribution.

## Declarations

### Ethics approval and consent to participate

Not applicable.

### Consent for publication

Not applicable.

### Availability of data and material

All methods and assessments described in this manuscript are publicly available at https://github.com/willtownes/quminorm-paper. The source code is licensed under LGPL-3.

### Authors’ contributions

FWT and RAI identified the problem. FWT proposed, derived, and implemented the quasi-UMI method. RAI provided guidance on refining the methods and evaluation strategies. FWT wrote the draft manuscript and revisions were suggested by RAI. All authors approved the final manuscript.

## Acknowledgements

The authors thank Martin Aryee, Jeff Miller, and Stephanie Hicks for valuable suggestions.

